# Robust scoring system to evaluate animal well-being in murine orthotopic breast cancer and ovarian cancer models

**DOI:** 10.64898/2025.12.18.695300

**Authors:** Ranya Al-Khaledy, Makiko Watanabe, Rebecca Shukla, Ladan Mashouri, Everett Stone, Mark Badeaux

## Abstract

Orthotopic models are critical to the development of new cancer therapies, as they have the greatest potential to mimic human cancer progression. However, orthotopic models are often underutilized due to practical challenges. These challenges include the need for specialized surgeries and the presence of complicating symptoms, such as ascites fluid accumulation in ovarian cancer and liver disease models, as well as ulceration of breast tumors growing in the mammary fat pad. An adaptable, quantitative animal health assessment system that utilizes simple, non-invasive, visual measurements can allow researchers to use these models to their full potential while safeguarding humane endpoints. The goal of this work was to establish such health scoring systems for translationally-relevant ovarian (ID8 Defb29-Vegf-A) and breast cancer (triple-negative model 4T1) models. We developed a tractable health assessment system by monitoring a combination of activity and responsiveness, coat condition, respiratory function, changes in body weight, posture, gait, tumor size (where measurable), and extent of ulceration, and scoring each of these parameters using a simple numerical scale. The scores at which euthanasia or other interventions were required were estimated and then adjusted based on clinical assessment by a team of researchers and veterinarians. These semi-quantitative health scoring systems were applied to survival experiments, in which mice were euthanized at a predetermined score or upon meeting a superseding health criterion, to confirm their applicability and effectiveness in maintaining humane endpoints. The scoring systems presented are intended to build on previously-established generic tumor model scoring systems that do not accommodate for the particular phenotypes present in orthotopic breast and ovarian cancer models.

## Introduction

Animal welfare is a central consideration in animal research. Scientific experiments involving animals are strictly regulated, primarily due to public concern and research ethics [1]. Since the Animal Welfare Act was passed in 1966, the concept of animal welfare has continually evolved as researchers’ understanding of research species has developed. Public opinion has driven stricter regulation by the Institutional Animal Care and Use Committees (IACUCs), boards comprised of researchers, veterinarians, and community members [2]. Researchers are required to submit proposals detailing the experimental objectives, benefits for society, and justification for the species and number of animals used in their work [2]. The purpose of these proposals is to enable IACUC committees to approve only experiments that have more potential for benefit than harm [3]. This type of required planning and justification has secondary benefits to researchers, such as reducing potentially wasteful experiments, and minimizing variables such as animal stress.

The development of anti-cancer therapies requires an expanded repertoire of orthotopic murine models that can recapitulate the microenvironment and natural disease course, including metastasis, of their human cancer equivalents [4]. Currently, there is great interest in the use of organoids to provide complex *in vitro* or *ex vivo* models, but while promising, these models have limitations that reduce their ability to faithfully mimic the tumor microenvironment, making the use of laboratory animals essential for progress. Mice are the most common living organism used in cancer research, and have been critical to the development and screening of cancer treatments [5, 6]. Murine models with clinically-relevant challenges such as ulceration, scabbing, or ascites fluid accumulation have increased pain and distress potential, and therefore often don’t fit within general health monitoring guidelines. These guidelines require analgesics or other methods to ameliorate malignancy-associated symptoms or pains, and humane termination of affected animals. However, the use of analgesics or premature termination of mice may compromise *in vivo* studies by shortening therapeutic windows and/or eliciting off-target effects that confound mechanistic studies; in total, this may prevent the research effort from producing scientifically-sound conclusions and translatable findings. To overcome this problem, a robust system that can monitor animals’ pain/stress level in an accurate and quantitative manner is required. Such a system will allow us to maintain humane endpoints while improving the quality of data on disease progression and standardization of experimental techniques [7, 8].

The 4T1 murine triple-negative breast cancer (TNBC) orthotopic model in BALB/c mice is one of only a handful of translationally-relevant syngeneic, orthotopic breast cancer models. Tumor cells implanted in the mammary fat pad rapidly and spontaneously metastasize to the liver, lungs, bone, brain and other distant sites, similar to human TNBC metastasis patterns [9–11]. The metastatic pattern and propensity of 4T1 tumors makes it a powerful model for studying the sequence of events and mechanisms at play in cancer metastasis, which is often neglected as a primary endpoint in preclinical models, despite being the major cause of cancer-related death in patients [12–15]. These tumors tend to ulcerate within two weeks of implantation due to the rapid growth of the tumors, leading to hypoxia, micro-hemorrhages, and necrosis [16–18]. Tumor ulceration can potentially lead to an increase in pain and distress to the animals, and is considered to be an endpoint for humane euthanasia. However, euthanizing mice as soon as they develop ulcers is not optimal for this model, as it significantly truncates the therapeutic window for testing anticancer therapies. In previous observations of 4T1 tumor-bearing mice, animals did not demonstrate behavioral indications of pain. However, as a prey species, mice are known to hide their pain and distress from potential predators. Therefore, close monitoring by personnel trained to recognize signs of pain and distress in mice allowed for meaningful data collection in experiments that continued past initial ulceration, while safeguarding the welfare of the mice, and maintaining ethical euthanasia criteria.

Orthotopic models of metastatic murine high-grade serous murine ovarian cancer (HGSOC) are established by intraperitoneal (i.p.) injection of ovarian cancer cell lines, e.g., the ID8 ovarian cancer cell line [19–22]. Surgical implantation directly into the ovarian bursa has been used, however it is invasive, stressful, and complicated [23, 24]. In contrast, i.p. injections place the cells directly into their preferred microenvironment, the peritoneal space, in a noninvasive, quick, and low-stress manner. ID8 and its genetically-modified versions are commonly used to model ovarian cancer progression in a complex environment [20]. ID8 tumor-bearing mice, similar to ovarian cancer patients, display significant ascites accumulation, presenting in mice as a rounding or swelling of the abdomen [21, 22, 25]. Assessment of disease progression is more difficult in this model compared to commonly-used subcutaneous models, as tumor dimensions cannot be measured directly with calipers, as is the standard with subcutaneous or other superficial tumors. Using genetically-modified cell lines that express luciferase or other markers allows for direct tumor measurement, but tumor growth has been observed to be affected by these highly immunogenic markers, and there are complications with signal dilution due to the extra abdominal fluid [22].

The use of analgesics is the most common way to treat animals in a state of pain and distress when euthanasia is not an option. However, mice with peritoneal ascites, as seen in ID8 tumor-bearing mice, still suffer from digestive, respiratory, and/or mobility issues, among other complications that analgesics cannot alleviate. In tumor models with early ulceration, such as 4T1, although analgesics can effectively alleviate pain, there is the potential for confounding effects of analgesic treatment on study endpoints. There are three common formats for analgesics; topical, NSAID, and opioid. Topical analgesics are not applicable for this type of pain relief, and there is some evidence suggesting the use of NSAIDs impact lung metastasis in 4T1 tumor-bearing mice even from a single dose [26]. Retroactive studies looking at opioid use in human cancers show that morphine and fentanyl can result in immunosuppression, and especially can alter T cell tumor penetration and effector versus regulatory T cell balance [27, 28]. These same studies observed that buprenorphine had little to no effect, but there has yet to be comparable data in mice. Additionally, most analgesic options are neither intended nor tested for extended use in mice. Rather, the intended use of analgesics is short-term use, usually for post-operative pain. Analgesic treatment from the time of scab formation to the time of euthanasia would require sustained treatment for two or more weeks. The lack of data regarding extended use of analgesics in mice is prohibitive, making a health scoring system that can detect animal distress with high sensitivity far more experimentally favorable.

Both 4T1 and ID8 orthotopic murine cancer cell lines represent translationally-relevant preclinical models that feature complications requiring exceptions to traditional euthanasia criteria in order to maximize their utility. Following traditional guidelines would limit the feasibility of the models as tools for preclinical anti-cancer therapeutic evaluation. Guideline exceptions can be justified with the use of a robust, sensitive monitoring system for pain and distress, ensuring that none of the animals are subject to undue stress, and preserving their quality of life.

## Materials and Methods

### Cell Culture

ID8 Defb29 Vegf-A_164_ (ID8-DV) cells were kindly provided by Dr. Jose R Conejo-Garcia and the Wistar Institute[21]. 4T1 cells were purchased from ATCC. Both cell lines were cultured in RPMI 1640 Medium (ATCC modification) with 10% fetal bovine serum and 1% Pen Strep. Cells were incubated at 37°C, 5% CO_2_, in T-75 flasks. Cells were passaged a minimum of 3 times after thawing, to ensure optimal and normal growth, and used for inoculation prior to 15 total passages. Cells in the logarithmic growth phase were trypsinized, centrifuged, and resuspended in RPMI 1640 serum-free medium. 4T1 cells were diluted to 5×10^5^ cells/mL in serum-free media to allow for 50 uL injections of 2.5×10^4^ cells. ID8-DV cells were diluted to 1×10^7^ cells/mL in PBS to allow for 200 uL injections of 2×10^6^ cells.

### Ethics statement

All animal work was approved by the University of Texas IACUC and conducted in accordance with recommendations from *The Guide for the Care and Use of Laboratory Animals*. The animal care and use program at the University of Texas at Austin is accredited by AAALAC, International. Developmental work was done under the supervision and in collaboration with the veterinarians on staff.

### Animals

Female C57BL/6J mice (8 to 12 weeks, JAX stock number 000664) and BALB/cJ mice (8 to 12 weeks, JAX stock number 000651) were obtained from The Jackson Laboratory (Bar Harbor, ME), and animals were acclimated for 1-2 weeks before tumor implantation. A total of 119 mice were used for the development of these health scoring systems: 60 mice to pilot and test the health scoring system for ID8-DV ovarian cancer and 59 to pilot and test the 4T1 breast cancer model.

Mouse colonies were monitored quarterly for adventitious viral, bacterial, protozoal, and parasitic pathogens by exhaust air dust testing. The exhaust air dust filter media were found to be negative for ectromelia virus, epizootic diarrhea of infant mice (mouse rotavirus), Hantann virus, lymphocytic choriomeningitis virus, LDH elevating virus, mouse adenovirus 1 and 2, mouse cytomegalovirus, mouse hepatitis virus, mouse norovirus, murine chapparvovirus, mouse parvovirus, mouse thymic virus, minute virus of mice, polyoma virus, pneumonia virus of mice, reovirus type 3, Sendai virus, Theiler’s meningoencephalitis virus, *Citrobacter rodentium, Clostridium piliforme, Corynebacterium bovis, Corynebacterium kutscheri, Filobacterium rodentium, Helicobacter* spp., *Klebsiella oxytoca, Klebsiella pneumoniae, Mycoplasma pulmonis, Pseudomonas aeruginosa, Rodentibacter heylii, Rodentibacter pneumotropicus, Salmonella* spp., *Staphylococcus aureus, Streptobacillus moniliformis, Streptococcus pneumoniae,* β-hemolytic *Streptococcus* (groups A, B, C, G), *Entamoeba muris, Giardia muris, Spironucleus muris, Tritrichomonas muris, Aspiculuris tetraptera, Syphacia muris, Syphacia obvelata, Myobia, Myocoptes,* and *Radfordia*.

### Housing and husbandry

Mice were housed at a density of 3 to 5 animals per cage under barrier conditions in autoclaved individually ventilated polysulfone cages with stainless steel wire-bar lids and filter tops (Blue Line Next 1285L; Tecniplast, West Chester, PA) at 60 air changes per hour, autoclaved 1/8” pelleted cellulose bedding (BioFresh, Patterson, NY), approximately 8 grams of autoclaved white crinkled paper strips (Enviro-dri; Shepherd Specialty Papers, Kalamazoo, MI), *ad libitum* access to *γ*-irradiated rodent diet (LabDiet 5053; PMI, St. Louis, MO), and *ad libitum* autoclaved domestic water provided in high temperature polycarbonate water bottles with stainless steel lids (Tecniplast, West Chester, PA). Caging components were changed at the following frequencies: Water bottles weekly, cages bottoms every 2 weeks, wire-bar lids and filter tops every 4 weeks. All cage and animal manipulations occurred within the housing room in a class II type A biological safety cabinet (BioGARD; The Baker Company, Sanford, ME), with the exception of anesthesia as described below. The housing room temperature and humidity were maintained at 72°F (± 4°F) and 30-70%, respectively, with fluorescent lighting in a 12:12h light:dark cycle (lights on at 0630, lights off at 1830). The room level disinfectant for surfaces and gloves was accelerated hydrogen peroxide 0.5% (Peroxigard; Virox Technologies, Oakville, ON Canada).

### Tumor implantation

Cell preparation and mammary fat pad injection of 4T1 cells was carried out according to published methods [16]. In short, one day or several hours before injection, depilatory cream was applied on the abdomen surrounding the left fourth nipple to remove fur for visualization of the MFP. To facilitate tumor cell injection, mice were anesthetized by isoflurane following the recommended General Anesthesia Regimen, with each mouse under isoflurane less than 10 minutes. Isoflurane was administered at 5% vol/vol for induction, and at 2% vol/vol via nose cone for the maintenance interval. Injection was performed by first tenting the skin around the MFP with forceps, piercing the skin with a 28-gauge needle, moving the needle just below the nipple, and pulling the needle up into the MFP [16]. After releasing the skin tent, 50 µL of 4T1 breast cancer cells were injected, and the needle was removed slowly with pressure applied to the injection site for 5 seconds after withdrawal by a cotton swab. For ID8-DV cells, 200 µL of cell suspension was injected intraperitoneally. The peritoneal space is the most biologically relevant location for ovarian cancer metastasis that does not require surgery to access [19].

### Monitoring of Mice

Researchers were not blinded as to mouse identity during assessments of animal health, but scoring of mice was confirmed to align with that of veterinary staff who were blinded. ID8-DV mice were inspected by visual assessment and weighed twice weekly until the onset of visible ascites or body weight gain of 10% relative to baseline. Once either of these metrics was reached, moistened food was placed on the cage prophylactically to alleviate cancer-related weight loss, and daily assessment was initiated. Assessment of 1) activity and responsiveness, 2) coat condition, 3) body posture, and 4) respiratory function was performed daily; body weight was assessed twice weekly until increasing to every other day after mice met a preset scoring criterion. The baseline weight was defined as the average of measurements taken between 0- and 14-days post-inoculation in order to allow for normal animal growth but prior to the onset of ascites-driven weight gain. Body weights of mice bearing ascites were measured daily only if they were near the criteria for ascites drainage at the previous observation.

Mice inoculated with breast cancer cells were checked by visual assessment, weighing, and measurement of tumors twice weekly until the onset of a pinprick size ulcer. At the onset of ulceration, all mice in the cage were observed daily, and scored weekly, for 1) activity and responsiveness, 2) coat condition, 3) gait, 4) (maximum) ulcer diameter, 5) body weight loss, and 6) tumor size. Tumors were measured using calipers and recording two diameters, one each along the x and y planes. The tumor size metric was scored based on the larger of the two diameters. Ulcer diameter was approximated with a caliper and only the apparent maximal diameter was measured. Once any mouse in a cage met a predetermined score, or if health deterioration was notable, scoring was performed every other day. Additionally, at this point the pulmonary assessment of advanced metastasis (PAAM) was included every other day, but not scored [29]. Representative images and additional guidelines for assessment of behavioral and health are provided in the supporting information.

### Development of the mouse well-being scoring systems

The initial scoring systems were based on the guidelines on humane endpoints, euthanasia, and analgesia provided by the University of Texas at Austin Office of Research Support and Compliance and the Standard Score Sheet for the Assessment of Wellbeing in Mice: Solid Tumour Models published by the University of Queensland Australia [30]. These initial templates were tweaked, incorporating strategies from several papers discussing novel scoring systems for specific applications, and guidelines for animals in cancer research were also consulted [31–34].

Pilot experiments were performed for both models to assess the tumor growth rate and ensure that the health scores aligned with veterinarians’ clinical assessment of the mice. Routine assessment was performed by biomedical researchers at the indicated frequency, with the addition of once or twice weekly meetings with the Animal Resource Center veterinarians to score mice and assess the appropriateness of actions triggered by the respective animals’ health score. In the case of the ID8-DV mice, an initial pilot study was conducted, with groups of 10 mice injected two weeks apart. After observation of ascites onset within the first cohort, the scoring system was updated and the improved system was tested with the second group. Two larger experiments of 20 mice each confirmed the system’s effectiveness and reproducibility. For 4T1 mice, a group of 10 mice was used to test first tumor implantation and growth rate. This was followed by a scoring system pilot with 24 mice. The pilot confirmed the primary tumor growth, metastatic, and ulceration rates. The health system was updated retroactively to eliminate categories that did not prove useful (body posture and respiratory effort) and replace them with the observable categories of gait and ulcer diameter. A final group of 25 mice was used to confirm the effectiveness and reproducibility of the health scoring system.

### Euthanasia

Euthanasia protocols were approved by The University of Texas at Austin IACUC. Mice meeting criteria for euthanasia were placed in a euthanasia chamber into which carbon dioxide was then introduced; secondary euthanasia, e.g., cervical dislocation, was performed when breathing ceased for several minutes. Euthanasia criteria were as follows: A total score of ≥10 for ID8-DV mice, 7 for 4T1 mice, or a severe score (3) in any individual category for either model. For ID8-DV mice meeting initial euthanasia criteria, amelioration of animal health was attempted by draining ascites fluid and observing the mouse for 12 to 24 hours post-drainage, after which time the mouse was re-scored. Mice were euthanized if they continued to display a score of 3 after ascites drainage. For both ID8-DV and 4T1 mice, a severe body weight change was only a euthanasia criterion if it persisted across two consecutive measurements. For 4T1 mice, there were three additional euthanasia criteria: failure of a PAAM [29], a tumor diameter exceeding 15 mm, or a maximum ulcer diameter exceeding 13 mm.

For ID8-DV mice, the overall body condition (level of hydration and normal fat deposits) was assessed with mice assigned an aggregate score of ≥6 [7, 8]. Due to the abdominal swelling in ascitic mice, this was simplified to an evaluation of the apparent feel of the spine while picking the mice up (scruffing). An increase in the prominence of the spine can indicate lean mass, or normal fat loss. If the spine was highly palpable, dehydration was assessed by a skin-pinch test; this entails tenting the skin at the back of the shoulders, and noting if the skin released back into place rapidly (hydrated) or slowly (dehydrated) [7]. In severe cases, weak or paralyzed rear legs may be observed [35]. Poor body condition was a euthanasia criterion if observed two days in a row.

### Ascites fluid, blood and tissue collection

Any mouse scoring over a 7 had its ascites drained using a 25-gauge needle attached to a 10 mL syringe. This was done by inserting the needle laterally to the bladder and parallel to the body, and slowly withdrawing fluid until it was no longer freely and easily flowing. The needle was first placed with the mouse head down, and then the body of the mouse was tilted to be lateral, enabling a freer flow of ascites fluid. The needle was stabilized by resting the syringe on the outer edge of the hand holding the mouse. The needle was removed from the syringe, dispensed into an EDTA tube, and placed on ice. Mice were weighed before and after drainage in order to collect data on lean body mass and cancer-related weight loss.

Blood was collected via cardiac puncture with a 25-gauge needle. The needle was then removed from syringe and blood was immediately dispensed into an EDTA tube and placed on ice. Lung tissue was collected as previously described [14], with the addition of a cardiac puncture prior to lung harvest to improve the visibility of the trachea, and the Miltenyi Biotech, Inc. Lung Dissociation Kit Mouse was used with the gentleMACS™ Octo Dissociator following the data sheet protocol for tissue digestion.

### Statistical Analysis

Statistical analysis was carried out using GraphPad Prism 10.4.1. Quantitative analyses were performed using a two-sided Student’s t-test, One-Way ANOVA, or the Mantel-Cox Survival. A p value of <0.05 was considered significant. For ID8 mice, a total of 19/60 mice were excluded from data analysis, including mice that did not develop tumors (n=15) and mice that were sacrificed prematurely to enable postmortem evaluation of tumor burden (n=4). All other animals were included in analyses, including all 59 4T1 mice.

## Results

### ID8-DV Ovarian Cancer

ID8-DV had a tumor take rate of 73% (44/60 mice). Mice that developed peritoneal ovarian cancer displayed a 10% weight gain, on average, 4 weeks after tumor cell inoculation (Fig 1A), due to ascites fluid buildup and in alignment with previously published data on this model [25]. Once weight gain reached an average of 10% above baseline for each group, daily health scoring was implemented (Fig 1B). The overall health score categories of activity and responsiveness, body posture, and body weight change were the biggest indicators of overall health. These were the most consistently altered criteria, and the earliest indicators of pain or distress after ascites accumulation. As health declined, coat condition and respiratory effort were also observably altered. The appearance of a rough coat and/or respiratory effort increase often aligned with a high enough score to require ascites drainage, a translationally-relevant intervention that relieves pain and fluid pressure-related organ dysfunction, e.g., respiratory difficulty, in ovarian cancer patients. Body weight loss below baseline was typically only evident following ascites drainage if the tap was complete, or when mice were at or near meeting a euthanasia criterion. Due to practical and animal health considerations, drainage of ascites was often incomplete. The majority of euthanized mice were found to have at least 1 mL of ascites fluid in their abdomens upon necropsy. Postmortem examination also revealed that mice presented with paler organs at the onset of ascites compared to healthy mice, but tumors were generally not grossly visible, and often were not apparent even with magnification. As the cancer progressed, small tumors were observed first and most easily along the peritoneum, diaphragm, and liver. With more advanced disease, a rough texture and/or opaque coating was evident across all organs in the peritoneal cavity (Fig S1). The presence of a significant amount of residual, often inaccessible fluid compromised the utility of body weight loss as a reliable marker of animal health, necessitating the use of weight *change* (gain or loss) as a scoring metric. Overall, upon ovarian cancer establishment, the health of each mouse was highly variable from day-to-day (Fig 1B), underscoring the need for a robust monitoring system.

**Figure 1:**
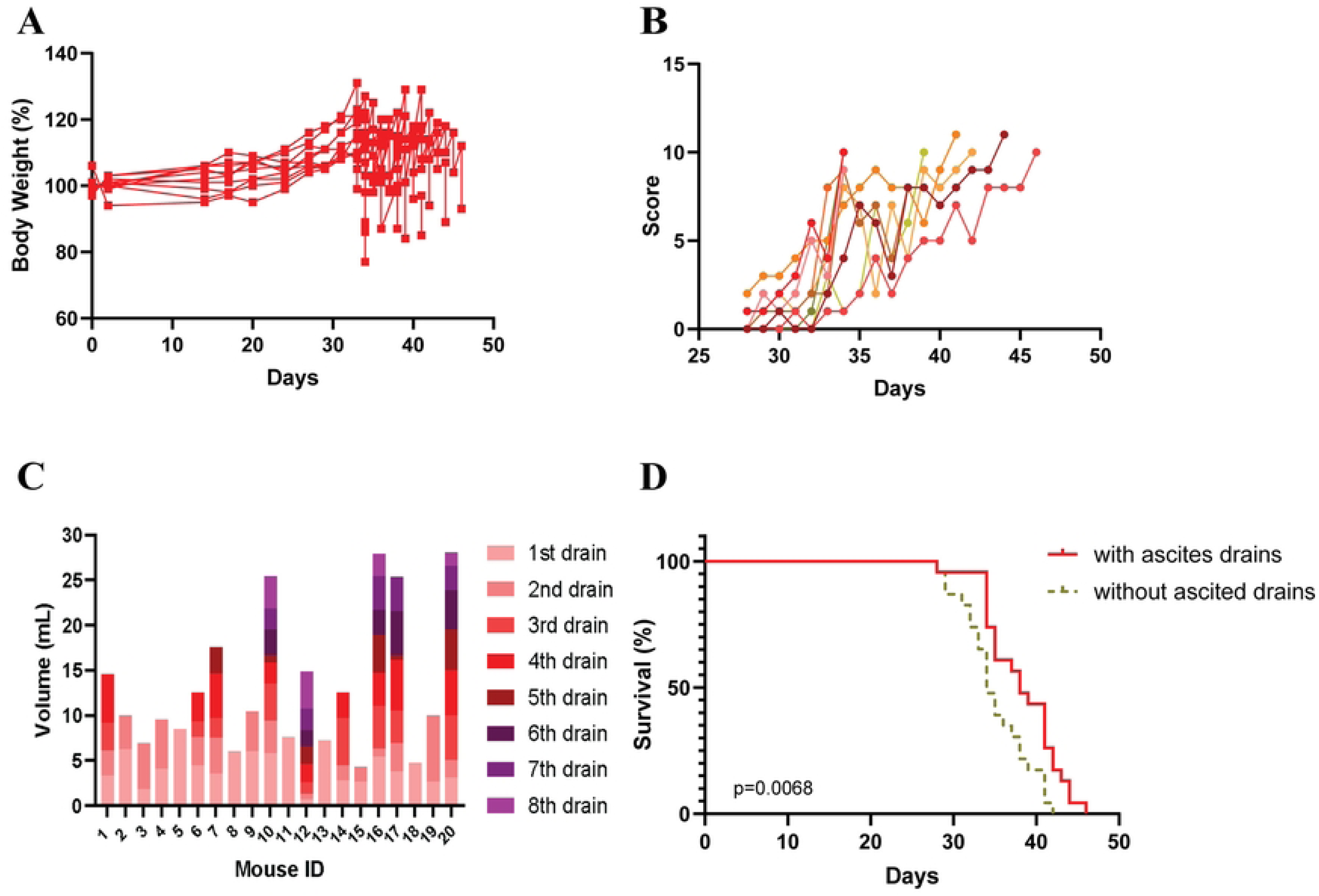
Scoring and health parameters of ovarian cancer-bearing mice. **A)** The individual weight fluctuation of ten mice, representative of all tumor inoculation cohorts. Each line represents a mouse exhibiting rapid weight gain and ascites drainage. Mice displayed an average 10% weight gain at week four. **B)** Data displays complete health scoring of the same 10 mice from **A**. Scoring began when weight change reached an average of 10% for the cage. **C)** Ascites was drained from mice with a health score >7 or weight change of >30%. **D)** Survival curves of mice from the 1^st^ (n=7), 2^nd^ (n=7), and 3^rd^ (n=6) tumor inoculation groups comparing euthanasia post-ascites withdrawal with theoretical survival without removal of ascites. All mice received PBS injections twice weekly. Mice euthanized before reaching euthanasia criteria were not included in the survival curves.

The presence of ascites fluid represented the most significant challenge in assessing animal health in the ID8DV model. The volume of fluid per ascites draw and time between fluid withdrawal was variable (Fig 1C). The number of drainages a single mouse received prior to euthanasia ranged from 1 to 8, with a median of 2 and a mean of 3.4 draws (Fig 1C). The average weight loss per ascites drainage was 3.6 g, but a single draw could be as little as 1 mL and as much as 10 mL. Draining ascites fluid when mice would otherwise be scored as terminally moribund and subsequently euthanized improved animal health scores; additionally, this intervention extended the survival of mice by roughly 4 days (Fig 1D). No mice were lost to spontaneous death, demonstrating the robustness of the scoring system in tracking the progressive decline in animal health. This is consistent with reports of ascites drainage extending animal survival [22], and underscores the utility of our health scoring system to measure the vitality of animals even in the presence of confounding factors such as ascites accumulation.

Ascites accumulation was the largest source of variability in this model; the time to first required ascites drainage ranged from 27 to 42 days post-tumor inoculation. The majority of animals had a higher red blood cell count in early ascites samples compared to ascites drawn closer to euthanasia (Fig 2A-C). Initially, ascites was only to be drawn when mice reached a weight threshold of 40% increase, but it was observed that mice with less than a 10% increase in weight occasionally had high scores and exhibited a highly-rounded appearance. We hypothesized that as ascites accumulated, the discomfort and mobility challenge resulted in lean mass loss masked by fluid accumulation. This was confirmed when lean weight was taken at necropsy, after draining the peritoneal space. In Fig 1A, the last point of each line is the weight after post-mortem total ascites drainage, and in most cases is markedly the lowest recorded body weight. Some mice appeared more comfortable with the extra abdominal fluid at the start, and most mice seemed to become accustomed to the ascites over time, as heavy and rounded mice scored only 4 or 5.

**Figure 2:**
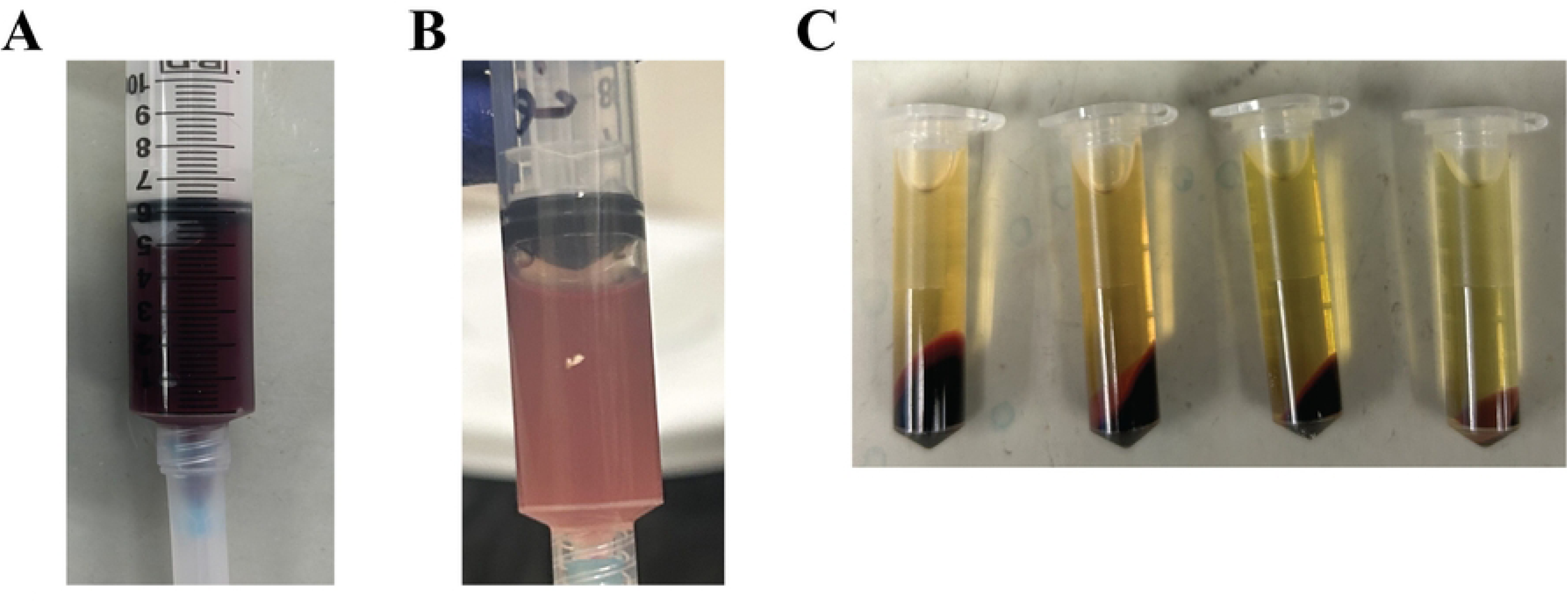
Ascites samples from ovarian cancer-bearing mice. **A)** A representative early (1^st^ or 2^nd^) ascites collection which appears bloody, **B)** a representative later (4^th^ - final) ascites collection which was a pale pink and cloudy, **C)** The supernatant of ascites, pale yellow in color, with varied red blood cell content as shown by the larger to smaller red blood cell pellets from left to right.

Two changes were made to the final rubric to accommodate for loss of lean mass: Moist chow or bacon softies were left on the cage floor for ease of access in order to mitigate lean mass loss starting when mice scored above a zero, and the criteria for ascites drainage was changed to either a weight gain of ≥30%, or a health score ≥7. After ascites removal, a rebound in the health score was usually observed, suggesting that this intervention improved animal health. However, ascites fluid regenerated rapidly, with mice gaining up to 5 grams of ascites-related weight in only one or two days. Mice whose health declined (as evident by health score increase) progressively following ascites drainage often, but not always, failed to display this rapid weight rebound and usually met euthanasia criteria the following day. Our system proved effective, aligned with the clinical assessment by the IACUC veterinarians, and prevented spontaneous death. The final scoring rubric is represented in Table 1. A comprehensive, one-page table with all scoring criteria, actions, and euthanasia criteria is available in Table S1.

**Table 1:** Clinical health scoring for ID8-DV mice. Mice were scored on a scale of Normal (0), Mild (1), Moderate (2), and Severe (3). Assessment included 1) activity and responsiveness, 2) coat condition,[35] 3) body posture,[36] 4) respiratory function, and 5) body weight change. Activity, responsiveness to stimulus, and coat condition were benchmarked at the start of the study; these parameters can be compared to healthy animals, if available. Respiratory function was scored as a relative increase or decrease in rate or effort compared to a healthy mouse. This was best observed by looking at the ribs and shoulders of the mouse. Actions were based on the sum of all categories’ score.

| Assessment | Score | Description |
| --- | --- | --- |
| Activity and responsiveness | 0 | Normal |
|  | 1 | Slightly less active compared to baseline |
|  | 2 | Marked reduction in activity OR isolation from cage mates |
|  | 3 | Non-responsive to nudging stimulus OR trembling when moving<br>OR stationary |
| Coat condition | 0 | Normal |
|  | 1 | Slightly rough coat |
|  | 2 | Intermittent rough coat |
|  | 3 | Coat is rough and unkempt throughout |
| Body posture | 0 | Normal |
|  | 1 | Slight hunching |
|  | 2 | Pronounced hunching |
|  | 3 | Severe hunching |
| Respiratory function | 0 | Normal |
|  | 1 | N/A |
|  | 2 | Slight increase in respiratory rate or effort relative to baseline |
|  | 3 | Marked increase in respiratory rate or effort relative to baseline |
| Body weight change | 0 | Gain of < 10% |
| | 1 | Gain of 10 to < 20% OR Loss of ≤ 5% |
|  | 2 | Gain of 20 to < 30% OR Loss of 5 to < 10% |
| | 3 | Gain of 30 to ≥ 40% OR Loss of 10 to ≥ 15% |

### 4T1 Breast Cancer

Overall, tumor growth and ulcer or scab formation was uniform among 4T1 tumor-bearing mouse cohorts. All mice inoculated with 4T1 cells developed a tumor. The terms “ulcer” and “scab” are used interchangeably for this study due to the dry, dark, scabbed appearance of all defects. Importantly, at no point was blood or any form of an open wound observed. Initial pinprick ulcers started forming on small tumors (∼200 mm^3^) less than two weeks from inoculation, and all mice bore some form of scab 3 weeks from inoculation (Fig 3A and B). Once a pinprick formed, scabs grew along with the tumor (Fig 3C).

**Figure 3:**
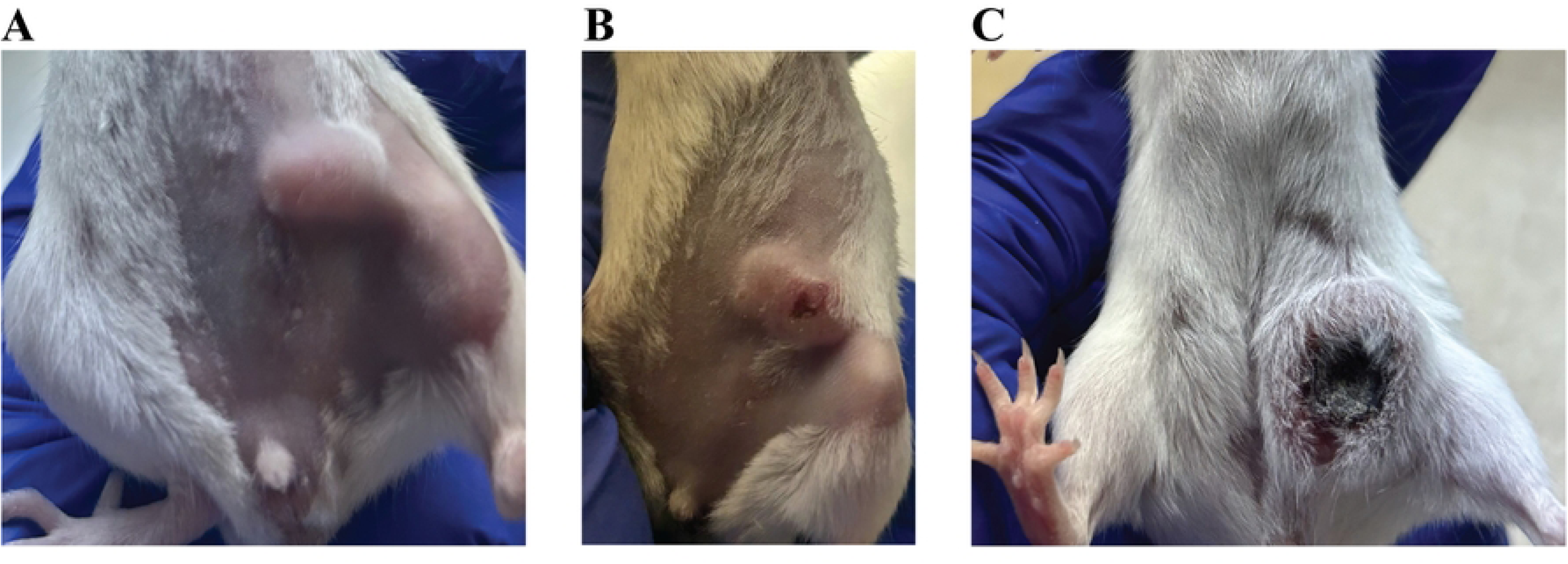
Ulceration patterns in orthotopic murine breast cancer. Mice bearing 4T1 tumors **A)** 17 days from tumor inoculation, prior to scab or ulcer formation, **B)** 17 days from tumor inoculation, with a pinprick ulcer present, and **C)** 30 days from tumor inoculation with a very late stage, large ulcer.

Primary tumor growth was the primary driver of increases in animal health score for the first 3 weeks of the experiment (Fig 4A and B). On the last day of survival, animal activity/responsiveness and coat condition were both observed to be altered, scoring above a zero. Mice noticeably did not display any observable changes with respect to body posture or respiratory effort. Therefore, both of these categories were eliminated from the final scoring rubric. There was no observable body weight loss throughout the experiment (Fig 4C), so food supplements were not included at any score threshold. Body weight was set to be measured only once weekly until a total score of ≥ 3, in order to balance the need for reliable quantitative assessments of health with the discomfort and stress the mice feel as a result of being handled by researchers. None of the mice displayed a failed PAAM, suggesting that euthanasia due to primary tumor size typically supersedes compromised lung function due to metastatic burden. PAAM was left in the health guide for mice with a score of ≥ 3, usually indicating a large tumor (between 10 and 15 mm in diameter), as an extra precaution due to the rapid metastasis common with 4T1 tumors [14, 15].

**Figure 4:**
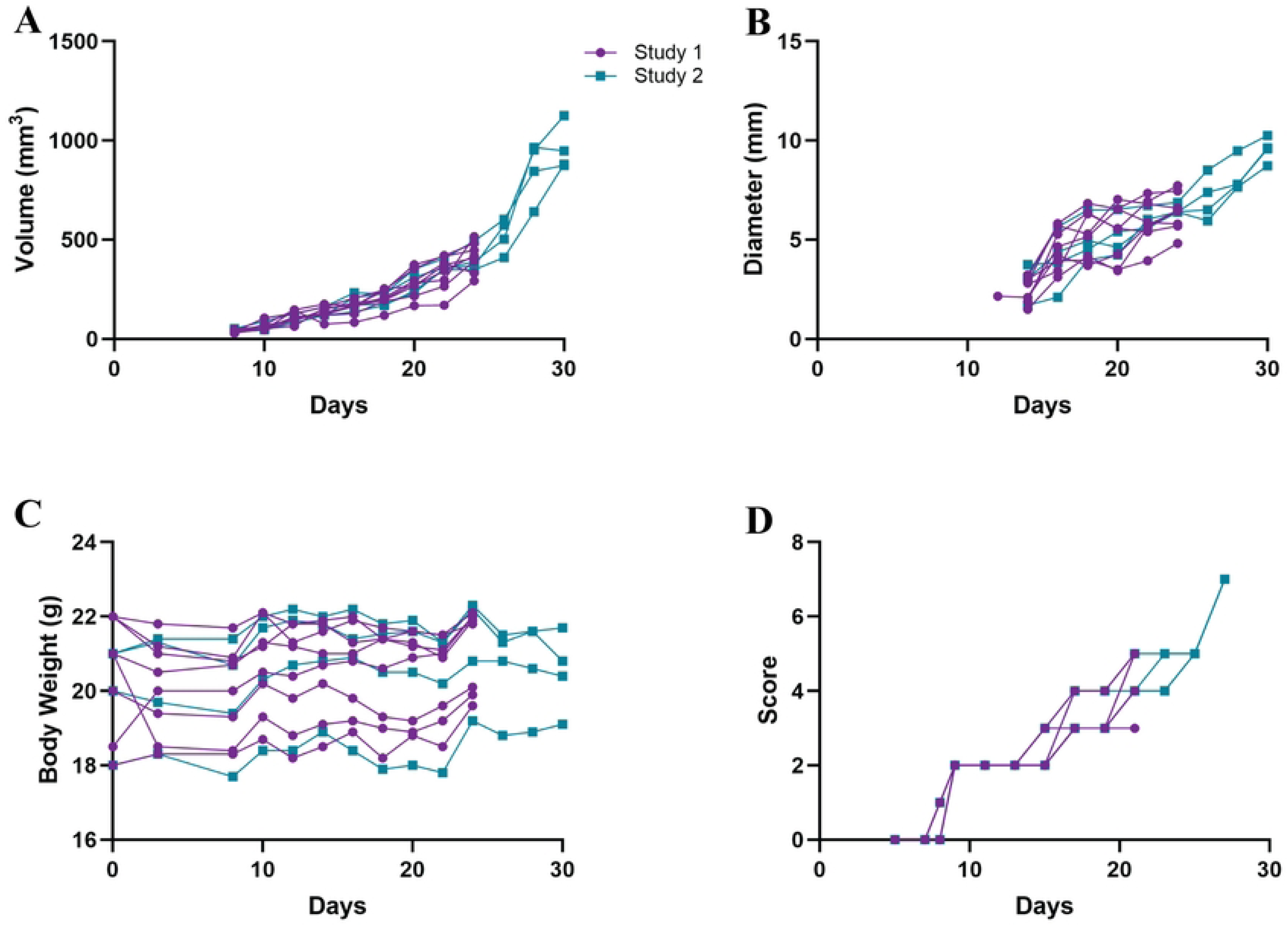
Monitoring of mice bearing 4T1 tumors. The 8 mice sacrificed at day 24 are shown in purple, while the 4 which were left until euthanasia criteria were met are in blue. Mice were given twice-weekly peritumoral injections of PBS. A) Primary tumor growth was measured by taking two diameters and multiplying 0.5 ∗ *L* ∗ *W*^2^, B) maximum ulcer diameter was taken as a single measurement, C) mouse body weight, and D) health score representing the sum of all six categories (Table 2).

**Table 2:** Clinical well-being scoring for 4T1 mice. Mice were scored on a scale of Normal (0), Mild (1), Moderate (2), and Severe (3). Activity, responsiveness to stimulus, gait, and coat condition were benchmarked at the start of the study; these parameters can be compared to healthy animals, if available. Actions were based on the sum of all categories’ score:

| Assessment | Score | Description |
| --- | --- | --- |
| Activity and<br>responsiveness | 0 | Normal |
|  | 1 | Slightly less active compared to baseline |
|  | 2 | Marked reduction in activity OR isolation from cage mates |
|  | 3 | Non-responsive to nudging stimulus OR trembling when moving OR stationary |
| Coat condition | 0 | Normal |
|  | 1 | Slightly rough coat |
|  | 2 | Intermittent rough coat |
|  | 3 | Coat is rough and unkempt throughout |
| Gait | 0 | Normal |
|  | 1 | Gait is affected in any way |
|  | 2 | N/A |
|  | 3 | N/A |
| Ulcer Diameter | 0 | Normal |
|  | 1 | < 6 mm maximum diameter |
|  | 2 | 6 to < 13 mm maximum diameter |
| | 3 | ≥ 13 mm maximum diameter |
| Body weight loss | 0 | < 5% |
|  | 1 | 5 to < 10% |
|  | 2 | 10 to < 15% |
| | 3 | ≥ 15% |
| Tumor size | 0 | < 5 mm in diameter |
|  | 1 | 5 to < 10 mm in diameter |
|  | 2 | 10 to < 15 mm in diameter |
| | 3 | ≥ 15 mm in diameter |

The gait of animals bearing large tumors was observed to be slightly altered, usually when tumor diameters reached between 9 and 10 mm (340 mm^3^). This threshold was reached approximately three weeks from tumor implantation. It is believed that the size and location of the tumor mechanically interfered with a natural gait. These tumors were located at the fourth left nipple, which is along the hip (Fig 3). Since no other signs of pain were observed when gait changed, this was not considered to be a sign of pain or distress. However, the observation allowed for an additional behavioral benchmark of increasing tumor size to be added to the assessment. Another metric was added after observing that ulcer diameter displayed a consistent growth pattern across mice. Therefore, in the 4T1 scoring rubric, the categories of body posture and respiratory effort were retroactively replaced with gait and ulcer diameter. The results of this scoring system are shown in Fig 4D.

Throughout the course of the experiments, there were no unexpected deaths, and tumor growth was consistent between tumor inoculations (time to ulceration, tumor volume, ulcer diameter) with slight deviations within groups. Table 2 below summarizes the final assessment rubric for mice bearing 4T1 tumors. A comprehensive, one-page table with all scoring criteria, actions, and euthanasia criteria is available in Table S2.

## Discussion

Publications providing specific insight into the methods of handling and caring for laboratory animals are becoming more common, as the ethical considerations for the use of animals evolve. IACUCs are also more frequently requiring these systems to be in place before approving planned experiments. There are publications providing guidelines to assess the health and well-being of mice in general or in the setting of cancer [7, 8, 34, 35]. However, many of these guidelines are more than 20 years old, and few provide robust suggestions on how to quantifiably assess animals or account for abnormal symptoms. There are also a handful of scoring systems for specific diseases [31–33, 37, 38], or protocols which detail thoroughly a specific assessment technique such as the PAAM ([29, 36]. Many of the scoring systems and technique assessments were published in the past 5 or 10 years, suggesting the field is moving towards using more comprehensive assessments in order to ensure animal welfare and collect better, more standardized data.

The weight gain, tumor growth, and survival with or without ascites drainage has been described for ID8 tumor-bearing mice [22]. However, even this work provided a minimal list of euthanasia criteria and no template for longitudinal tracking of these metrics. Similarly, in 4T1 tumor-bearing mice, the presence of a scab or ulcer is a regularly-observed phenomenon, and is noted in many publications [10, 15, 17, 18, 39]. While it is easy to find mention of this phenomena, most publications provide very little or no information about steps taken to align with animal welfare guidelines while continuing to treat and observe mice bearing scabbed or ulcerated tumors, despite this typically being considered a euthanasia criterion [7, 30]. The vast majority of the literature simply states that all protocols are approved by the IACUC committee, with no references or details about the protocol.

We present here a scoring system for robust monitoring of animal health in the context of several different tumor types. From a practical perspective, our health assessment framework is able to prevent spontaneous animal death, as the gradual decline in animal health can be used to accurately predict mortality and intervene judiciously with euthanasia. This can be done even in the presence of complicating factors such as ascites accumulation. By employing this system or a similar framework, it is possible to extend the therapeutic window in models of ovarian cancer and limit loss of animals to unforeseen mortality.

Translational relevance is a concern for any animal model, particularly mouse models of cancer, which have generally shown a high level of discordance with outcomes in human patients. Therefore, evaluating how murine models align with the disease state in humans is important for understanding the strengths and shortcomings of a given model. Importantly, our scoring system facilitates translationally-relevant therapeutic interventions, including ascites drainage, which is a common practice that relieves pain and distress in ovarian cancer patients. Murine models of ovarian cancer deviate from human ovarian cancer progression in several respects. First, ascites accumulation in mice seems to occur more consistently and rapidly in mice compared to humans [24, 40, 41]. The total number of drains and the cumulative volume of ascites drained from a mouse correlated positively with survival in mice bearing ID8-DV tumors, in opposition to the poorer prognosis in humans with greater speed and volume of ascites accumulation [24]. More closely following common practices for human patient ascites drainage may improve the clinical relevance of the model. Additionally, in humans, the appearance of ascites fluid is described as pale yellow [41]. In mice, the ascites fluid collected ranged from a deep red to a very pale, cloudy pink. After red blood cell removal, the fluid supernatant was typically a pale, clear yellow [42]. None of the ascites collected from mice could be described as pale yellow at the time of collection, suggesting a fundamental difference in cellular and metabolic composition in ovarian cancer-induced ascites between humans and mice. Comparing the cellular and molecular composition of human and murine ascites would enable a better understanding of the potential limitations of the model as a tool for modeling human pathology.

One of the notable differences in the presentation of 4T1 tumors compared to human breast cancer is the consistent ulceration of 4T1 tumors. In the mouse model, a small pinprick is first observed very near or on the nipple, and this lesion grows along with the tumor [17, 39]. In comparison, a relatively small proportion of human patients experience changes in their nipple appearance, but this is mild compared to what is observed in 4T1 tumors [43]. Additionally, it has been observed that the scabs may present challenges to imaging, complicate models, and have less blood flow compared to tumors without major scab or ulceration issues [10, 17, 18]. A major benefit of 4T1 tumors is the spontaneous metastasis to the lungs and other organs. This makes it a powerful tool for the study of both TNBC tumors and metastasis. 4T1 cells can be quantitatively detected in tissues [14] or as circulating tumor cells via luminescence [44]. In this case, we determined that by 24 days from tumor implantation, lung metastasis was easily detectable via colony analysis using published methods (data not shown) [14].

Ideally, the use of these systems will be expanded to encompass models with similar complications. The ovarian cancer rubric could easily be incorporated for alternative ovarian cancer cell lines [20, 25, 45], as well as models of pancreatic, colorectal, liver, and endometrial cancers which also cause ascites. Murine cirrhosis and liver disease models could potentially utilize a similar rubric since ascites is a common symptom [38, 46]. Tumor size and ulceration are striking visual phenomena, but often do not directly reflect the pain and quality of life of a given animal. Therefore, incorporating these parameters into a health scoring system alongside other measures of animal wellbeing ensures their quality of life in the experimental setting. In the future, tumors that feature similar dry, consistently-scabbed lesions could be considered for longer treatment windows past the point of initial ulceration, so long as animal health and quality of life is maintained. Finally, our scoring system, or a derivative thereof, should have utility in metastasis models where the primary tumor is resected, as it is based on generic behavioral and physical signs of pain and distress, and secondary aspects of the scoring system, such as the PAAM, may be of greater importance.

The scoring systems provided here are highly adaptable, and would require only additional small-scale pilot experiments to evaluate their efficacy, reproducibility, and need for additional metrics. The adaption of these rubrics to similar models would save time in early development and enable a well-planned pilot experiment with fewer animals compared to starting with less-applicable rubrics or designing a system *de* novo. Ideally, the availability of these types of scoring systems will also lower the barrier for researchers to begin using 4T1 or ID8 and their derivative cell lines. It is important that collaboration with veterinary staff is present when adapting these models, as the alignment of a veterinary professionals’ clinical assessment with the score and euthanasia is perhaps the most critical aspect of using a well-being scoring system effectively.

Publicly-available mouse behavioral or health scoring systems and guidelines can enable the widespread use of complicated models [34]. It is crucial that the proposed rubrics align with the veterinary staffs’ professional assessment of health and required euthanasia of animals. Both rubrics developed and described here were in alignment with clinical assessments and approved for use by the University of Texas at Austin’s Institutional Care and Use Committee. The proposed well-being scoring systems and accompanying rubrics can help empower researchers to incorporate both ID8-DV and 4T1 for the study of female cancers in their research. Furthermore, both of these rubrics could be expanded to encompass models with similar complications.

## Acknowledgments

The authors wish to thank Dr. George Georgiou for his thoughtful comments and support of this research. The authors would also like to thank Dr. Jose R Conejo-Garcia and the Wistar Institute for their generous gift of the ID8 Defb29 Vegf-A_164_ cell line.

## Supporting Information

**Table S1: *Mouse Health Scoring Systems for Ovarian Cancer Models.*** When mouse body weight is >30% above baseline or when their total score is >= 7, drain ascites. *Up to ten drainages are allowed prior to euthanasia*.

**Table S2: *Mouse Health Scoring Systems for Breast Cancer Models*** Scoring for wellbeing will be performed relative to the following assessment criteria. Any mouse with an ulcer needs to be observed daily. Scoring can be done once weekly unless a mouse displays signs of deterioration, at which point scoring should be every other day.

---

**Figure S1:**
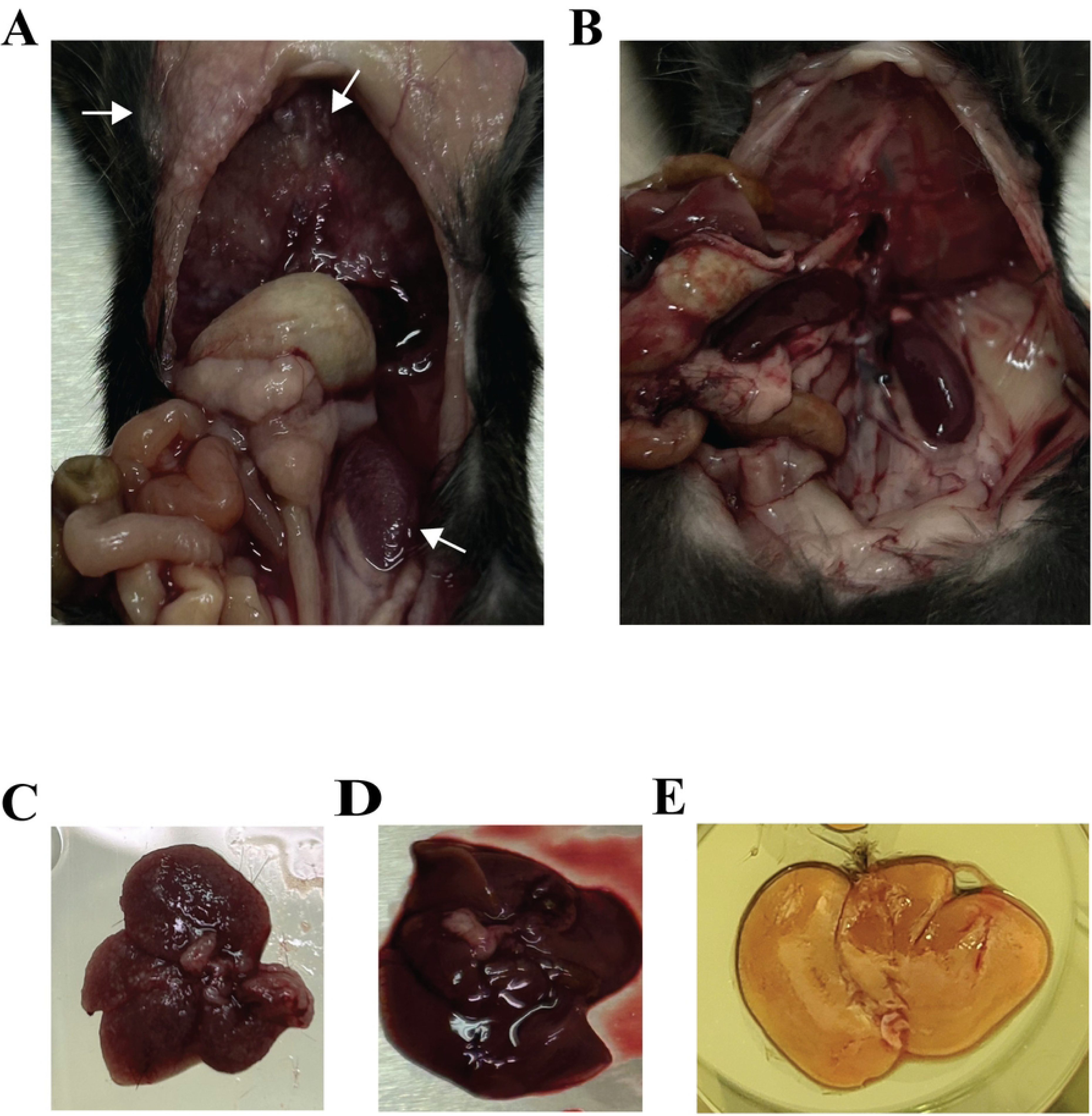
With more advanced disease, organs appeared to have a rough and/or opaque coating across the majority of organs in the peritoneal cavity. **A)** Red arrows are pointing to this rough texture which are small metastases coating the peritoneum, diaphragm, and kidney most notably. **B)** A representative non-cancerous mouse where the diaphragm, kidneys, and peritoneum appear more vascularized (deep red), and with a smooth shiny texture. **C)** A late-stage ovarian cancer liver with a rough opaque metastatic coating. **D)** An early-stage ovarian cancer mouse. This liver does not appear rough textured with metastases, but appeared paler or bleached and the mouse had ascites. **E)** A healthy liver with a shiny texture and dark red color.

